# Statistical shape analysis of large datasets based on diffeomorphic iterative centroids

**DOI:** 10.1101/363861

**Authors:** Claire Cury, Joan A. Glaunès, Roberto Toro, Marie Chupin, Gunter Shumann, Vincent Frouin, Jean-Baptiste Poline, Olivier Colliot, and the Consortium Imagen

## Abstract

In this paper, we propose an approach for template-based shape analysis of large datasets, using diffeomorphic centroids as atlas shapes. Diffeomorphic centroid methods fit in the Large Deformation Diffeomorphic Metric Mapping (LDDMM) framework and use kernel metrics on currents to quantify surface dissimilarities. The statistical analysis is based on a Kernel Principal Component Analysis (Kernel PCA) performed on the set of momentum vectors which parametrize the deformations. We tested the approach on different datasets of hippocampal shapes extracted from brain magnetic resonance imaging (MRI), compared three different centroid methods and a variational template estimation. The largest dataset is composed of 1000 surfaces, and we are able to analyse this dataset in 26 hours using a diffeomorphic centroid. Our experiments demonstrate that computing diffeomorphic centroids in place of standard variational templates leads to similar shape analysis results and saves around 70% of computation time. Furthermore, the approach is able to adequately capture the variability of hippocampal shapes with a reasonable number of dimensions, and to predict anatomical features of the hippocampus in healthy subjects.

## 1 Introduction

Statistical shape analysis methods are increasingly used in neuroscience and clinical research. Their applications include the study of correlations between anatomical structures and genetic or cognitive parameters, as well as the detection of alterations associated with neurological disorders. A current challenge for methodological research is to perform statistical analysis on large databases, which are needed to improve the statistical power of neuroscience studies.

A common approach in shape analysis is to analyse the deformations that map individuals to an atlas or template, e.g. (1)(2)(3)(4)(5). The three main components of these approaches are the underlying deformation model, the template estimation method and the statistical analysis itself. The Large Deformation Diffeomorphic Metric Mapping (LDDMM) framework (6)(7)(8) provides a natural setting for quantifying deformations between shapes or images. This framework provides diffeomorphic transformations which preserve the topology and also provides a metric between shapes. The LDDMM framework is also a natural setting for estimating templates from a population of shapes, because such templates can be defined as means in the induced shape space. Various methods have been proposed to estimate templates of a given population using the LDDMM framework (4)(9)(10)(3). All methods are computationally expensive due the complexity of the deformation model. This is a limitation for the study of large databases.

In this paper, we present a fast approach for template-based statistical analysis of large datasets in the LDDMM setting, and apply it to a population of 1000 hippocampal shapes. The template estimation is based on diffeomorphic centroid approaches, which were introduced at the Geometric Science of Information conference GSI13 (11,12). The main idea of these methods is to iteratively update a centroid shape by successive matchings to the different subjects. This procedure involves a limited number of matchings and thus quickly provides a template estimation of the population. We previously showed that these centroids can be used to initialize a variational template estimation procedure (12), and that even if the ordering of the subject along itarations does affect the final result, all centres are very similar. Here, we propose to use these centroid estimations directly for template-based statistical shape analysis. The analysis is done on the tangent space to the template shape, either directly through Kernel Principal Component Analysis (Kernel PCA (13)) or to approximate distances between subjects. We perform a thorough evaluation of the approach using three datasets: one synthetic dataset and two real datasets composed of 50 and 1000 subjects respectively. In particular, we study extensively the impact of different centroids on statistical analysis, and compare the results to those obtained using a standard variational template method. We will also use the large database to predict, using the shape parameters extracted from a centroid estimation of the population, some anatomical variations of the hippocampus in the normal population, called Incomplete Hippocampal Inversions and present in 17% of the normal population (14). IHI are also present in temporal lobe epilepsy with a frequency around 50% (15), and is also involved in major depression disorders (16).

The paper is organized as follows. We first present in section 2 the mathematical frameworks of diffeomorphisms and currents, on which the approach is based, and then introduce the diffeomorphic centroid methods in section 3. Section 4 presents the statistical analysis. The experimental evaluation of the method is then presented in Section 5.

## 2 Mathematical Frameworks

Our approach is based on two mathematical frameworks which we will recall in this section. The Large Deformation Diffeomorphic Metric Mapping framework is used to generate optimal matchings and quantify differences between shapes. Shapes themselves are modelled using the framework of currents which does not assume point-to-point correspondences and allows performing linear operations on shapes.

### 2.1 LDDMM framework

Here we very briefly recall the main properties of the LDDMM setting. See (6, 7, 8) for more details. The Large Deformation Diffeomorphic Metric Mapping framework allows analysing shape variability of a population using diffeomorphic transformations of the ambient 3D space. It also provides a shape space representation which means that shapes of the population are seen as points in an infinite dimensional smooth manifold, providing a continuum between shapes.

In the LDDMM framework, deformation maps *φ*: ℝ^3^ → ℝ^3^ are generated by integration of time-dependent vector fields *v*(*x*,⋅) with *x* ∈ ℝ^3^ and *t* ∈ [0, 1]. If *v*(*x*, *t*) is regular enough, i.e. if we consider the vector fields (*v*(⋅, *t*))_*t*∈[0,1]_ in *L*^2^([0, 1], *V*), where *V* is a Reproducing Kernel Hilbert Space (RKHS) embedded in the space of *C*^1^ vector fields vanishing at infinity, then the transport equation:

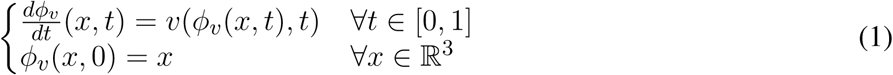

has a unique solution, and one sets *φ_v_* = *ϕ_v_*(⋅, 1) the diffeomorphism induced by *v*(*x*, ⋅). The induced set of diffeomorphisms 𝓐_*V*_ is a subgroup of the group of *C*^1^ diffeomorphisms. The regularity of velocity fields is controlled by:

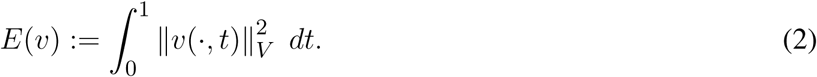

The subgroup of diffeomorphisms 𝓐_*V*_ is equipped with a right-invariant metric defined by the rules: ∀*φ*, *ψ* ∈ 𝓐_*V*_,

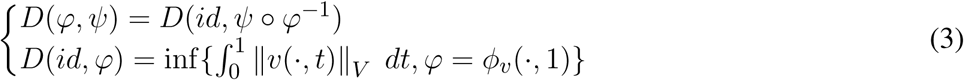

i.e. the infimum is taken over all *v* ∈ *L*^2^([0, 1], *V*) such that *φ_v_* = *φ*. *D*(*φ*, *ψ*) represents the shortest length of paths connecting *φ* to *ψ* in the diffeomorphisms group.

### 2.2 Momentum vectors

In a discrete setting, when the matching criterion depends only on *φ_v_* via the images *φ_v_*(*x_p_*) of a finite number of points *x_p_*(such as the vertices of a mesh) one can show that the vector fields *v*(*x*, *t*) which induce the optimal deformation map can be written via a convolution formula over the surface involving the reproducing kernel *K_V_* of the RKHS *V*:

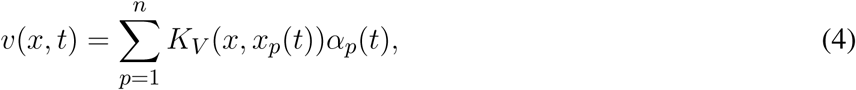

where *x_p_*(*t*) = *ϕ_v_*(*x_p_*, *t*) are the trajectories of points *x_p_*, and *α_p_*(*t*) ∈ ℝ^3^ are time-dependent vectors called momentum vectors, which completely parametrize the deformation. Trajectories *x_p_*(*t*) depend only on these vectors as solutions of the following system of ordinary differential equations:

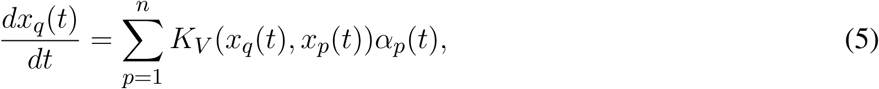

for 1 ≤ *q* ≤ *n*. This is obtained by plugging formula 4 for the optimal velocity fields into the flow equation 1 taken at *x* = *x_q_*. Moreover, the norm of *v*(⋅, *t*) also takes an explicit form:

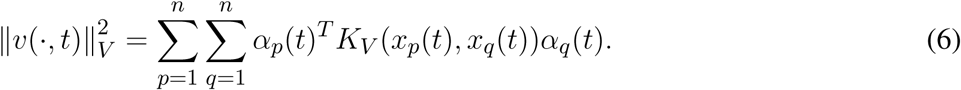

Note that since *V* is a space of vector fields, its kernel *K_V_*(*x*, *y*) is in fact a 3×3 matrix for every *x*, *y* ∈ ℝ^3^. However we will only consider scalar invariant kernels of the form 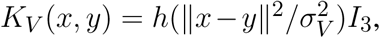 where *h* is a real function (in our case we use the Cauchy kernel *h*(*r*) = 1/(1 + *r*)), and *σ_V_* a scale factor. In the following we will use a compact representation for kernels and vectors. For example equation 6 can be written:

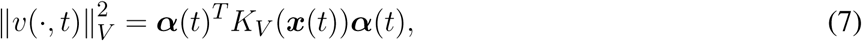

where ***α***(*t*) = (*α_p_*(*t*))_p=1…*n*_, ∈ ℝ^3×*n*^, ***x***(*t*) = (*x_p_*(*t*))_p=1…*n*_, ∈ ℝ^3×*n*^ and *K_V_*(***x***(*t*)) the matrix of *K_V_* (*x_p_*(*t*), *x_q_*(*t*)).

#### Geodesic shooting

The minimization of the energy *E*(*v*) in matching problems can be interpreted as the estimation of a length-minimizing path in the group of diffeomorphisms 𝓐_*V*_, and also additionally as a length-minimizing path in the space of point sets when considering discrete problems. Such length-minimizing paths obey geodesic equations (see (3)) which write as follows:

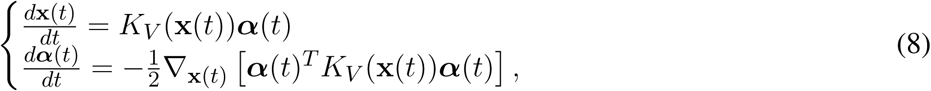

Note that the first equation is nothing more than equation 5 which allows to compute trajectories *x_p_*(*t*) from any time-dependent momentum vectors *α_p_*(*t*), while the second equation gives the evolution of the momentum vectors themselves. This new set of ODEs can be solved from any initial conditions (*x_p_*(0), *α_p_*(0)), which means that the initial momentum vectors *α_p_*(0) fully determine the subsequent time evolution of the system (since the *x_p_*(0) are fixed points). As a consequence, these initial momentum vectors encode all information of the optimal diffeomorphism. For example, the distance *D*(*id*, *φ*) satisfies

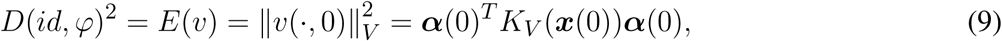

We can also use geodesic shooting from initial conditions (*x_p_*(0), *α_p_*(0)) in order to generate any arbitrary deformation of a shape in the shape space.

### 2.3 Shape representation: Currents

The use of currents ((17, 18)) in computational anatomy was introduced by J. Glaunés and M. Vaillant in 2005 (19)(20) and subsequently developed by Durrleman ((21)). The basic idea is to represent surfaces as currents, i.e. linear functionals on the space of differential forms and to use kernel norms on the dual space to express dissimilarities between shapes. Using currents to represent surfaces has some benefits. First it avoids the point correspondence issue: one does not need to define pairs of corresponding points between two surfaces to evaluate their spatial proximity. Moreover, metrics on currents are robust to different samplings and topological artefacts and take into account local orientations of the shapes. Another important benefit is that this model embeds shapes into a linear space (the space of all currents), which allows considering linear combinations such as means of shapes in the space of currents.

Let us briefly recall this setting. For sake of simplicity we present currents as linear forms acting on vector fields rather than differential forms which are an equivalent formulation in our case. Let *S* be an oriented compact surface, possibly with boundary. Any smooth vector field *w* of ℝ^3^ can be integrated over *S* via the rule:

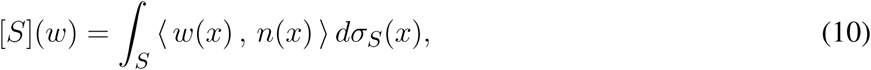

with *n*(*x*) the unit normal vector to the surface, *dσ_s_* the Lebesgue measure on the surface *S*, and [*S*] is called a 2-current associated to *S*.

Given an appropriate Hilbert space (*W*, 〈⋅, ⋅〉_*W*_) of vector fields, continuously embedded in 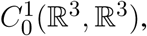 the space of currents we consider is the space of continuous linear forms on *W*, i.e. the dual space *W*^*^. For any point *x* ∈ ℝ^3^ and vector *α* ∈ ℝ^3^ one can consider the Dirac functional 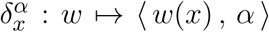 which belongs to *W*^*^. The Riesz representation theorem states that there exists a unique *u* ∈ *W* such that for all 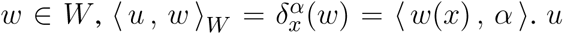 is thus a vector field which depends on *x* and linearly on *α*, and we write it *u* = *K_W_*(⋅, *x*)*α*. *K_W_*(*x*, *y*) is a 3 × 3 matrix, and *K_W_*: ℝ^3^ × ℝ^3^ → ℝ^3×3^ the mapping called the reproducing kernel of the space *W*. Thus we have the rule

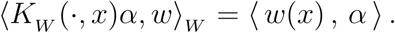

Moreover, applying this formula to *w* = *K_W_* (⋅, *y*)*β* for any other point *y* ∈ ℝ^3^ and vector *β* ∈ ℝ^3^, we get

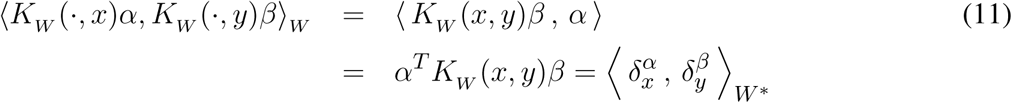

Using equation 11, one can prove that for two surfaces *S* and *T*,

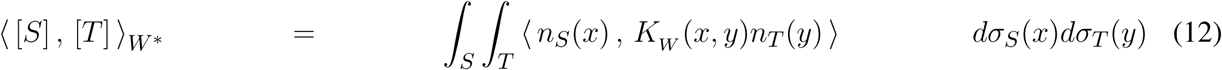

This formula defines the metric we use as data attachment term for comparing surfaces. More precisely, the difference between two surfaces is evaluated via the formula:

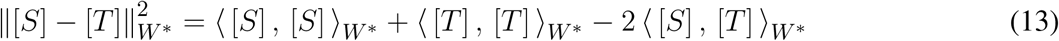

The type of kernel fully determines the metric and therefore will have a direct impact on the behaviour of the algorithms. We use scalar invariant kernels of the form 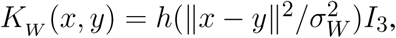 where *h* is a real function (in our case we use the Cauchy kernel *h*(*r*) = 1/(1 + *r*)), and *σ_W_* a scale factor.

Note that the varifold (22) can be also use for shape representation without impacting the methodology. The shapes we used for this study are well represented by currents.

### 2.4 Surface matchings

We can now define the optimal match between two currents [*S*] and [*T*], which is the diffeomorphism minimizing the functional

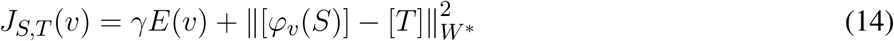

This functional is non convex and in practice we use a gradient descent algorithm to perform the optimization, which cannot guarantee to reach a global minimum. We observed empirically that local minima can be avoided by using a multi-scale approach in which several optimization steps are performed with decreasing values of the width *σ_W_* of the kernel *K_W_* (each step provides an initial guess for the next one). Evaluations of the functional and its gradient require numerical integrations of high-dimensional ordinary differential equations (see equation 5), which is done using Euler trapezoidal rule. Note that three important parameters control the matching process: *γ* controls the regularity of the map, *σ_V_* controls the scale in the space of deformations and *σ_W_* controls the scale in the space of currents.

### 2.5 GPU implementation

To speed up the matchings computation of all methodes used in this study (the variational template and the different centroid estimation algorithms), we use a GPU implementation for the computation of kernel convolutions. This computation constitutes the most time-consuming part of LDDMM methods. Computations were performed on a Nvidia Tesla C1060 card. The GPU implementation can be found here: http://www.mi.parisdescartes.fr/~glaunes/measmatch/measmatch040816.zip

## 3 Diffeomorphic Centroids

Computing a template in the LDDMM framework can be highly time consuming, taking a few days or some weeks for large real-world databases. Here we propose a fast approach which provides a centroid correctly centred among the population.

### 3.1 General idea

The LDDMM framework, in an ideal setting (exact matching between shapes), sets the template estimation problem as a centroid computation on a Riemannian manifold. The Fréchet mean is the standard way for defining such a centroid and provides the basic inspiration of all LDDMM template estimation methods.

If *x^i^*, 1 ≤ *i* ≤ *N* are points in ℝ^*d*^, then their centroid is defined as

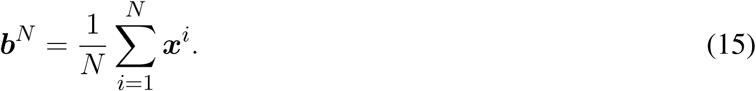

It also satisfies the following two alternative characterizations:

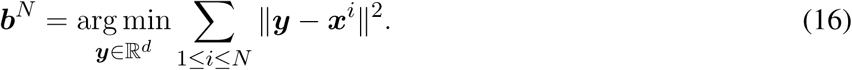

and

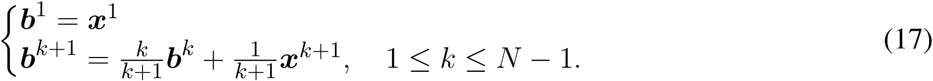

Now, when considering points *x^i^* living on a Riemannian manifold *M*(we assume *M* is path-connected and geodesically complete), the definition of ***b**^N^* cannot be used because *M* is not a vector space. However the variational characterization of ***b**^N^* as well as the iterative characterization, both have analogues in the Riemannian case. The Fréchet mean is defined under some hypotheses (see (23)) on the relative locations of points *x^i^* in the manifold:

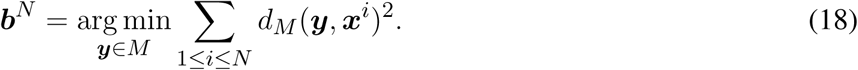

Many mathematical studies (as for example Kendall (24), Karcher (25) Le (26), Afsari (27, 28), Arnaudon (23)), have focused on proving the existence and uniqueness of the mean, as well as proposing algorithms to compute it. However, these approaches are computationally expensive, in particular in high dimension and when considering non trivial metrics. An alternative idea consists in using the Riemannian analogue of the second characterization:

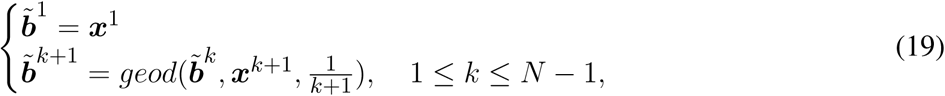

where *geod*(***y***, ***x***, *t*) is the point located along the geodesic from ***y*** to ***x***, at a distance from ***y*** equal to *t* times the length of the geodesic. This does not define the same point as the Fréchet mean, and moreover the result depends on the ordering of the points. In fact, all procedures that are based on decomposing the Euclidean equality 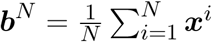 as a sequence of pairwise convex combinations lead to possible alternative definitions of centroid in a Riemannian setting. However, this should lead to a fast estimation. We hypothesize that, in the case of shape analysis, it could be sufficient for subsequent template based statistical analysis. Moreover, this procedure has the side benefit that at each step ***b**^k^* is the centroid of the ***x**^i^*, 1 ≤ *i* ≤ *k*.

In the following, we present three algorithms that build on this idea. The two first methods are iterative, and the third one is recursive, but also based on pairwise matchings of shapes.

### 3.2 Direct Iterative Centroid (IC1)

The first algorithm roughly consists in applying the following procedure: given a collection of *N* shapes *S_i_*, we successively update the centroid by matching it to the next shape and moving along the geodesic flow. More precisely, we start from the first surface *S*_1_, match it to *S*_2_ and set *B*_2_ = *ϕ*_*v*^1^_(*S*_1_, 1/2). *B_2_* represents the centroid of the first two shapes, then we match *B*_2_ to *S*_3_, and set as *B*_3_ = *ϕ*_*v*^1^_(*B*_2_, 1/3). Then we iterate this process (see Algorithm 1).

#### Algorithm 1: Iterative Centroid 1 (IC1)

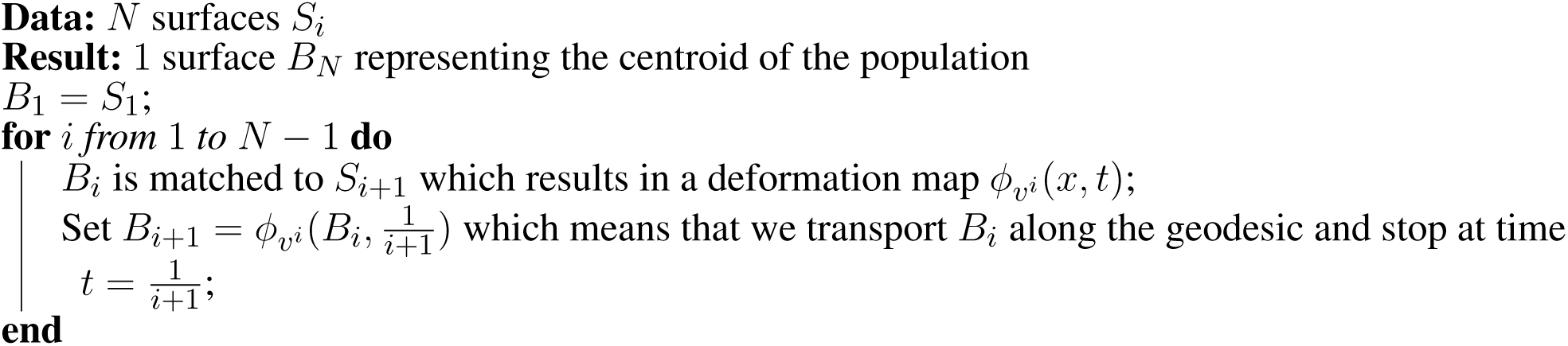

### 3.3 Centroid with averaging in the space of currents (IC2)

Because matchings are not exact, the centroid computed with the IC1 method accumulates small errors which can have an impact on the final centroid. Furthermore, the final centroid is in fact a deformation of the first shape *S*_1_, which makes the procedure even more dependent on the ordering of subjects than it would be in an ideal exact matching setting. In this second algorithm, we modify the updating step by computing a mean in the space of currents between the deformation of the current centroid and the backward flow of the current shape being matched. Hence the computed centroid is not a surface but a combination of surfaces, as in the template estimation method. The algorithm proceeds as presented in Algorithm 2.

#### Algorithm 2: Iterative Centroid 2 (IC2)

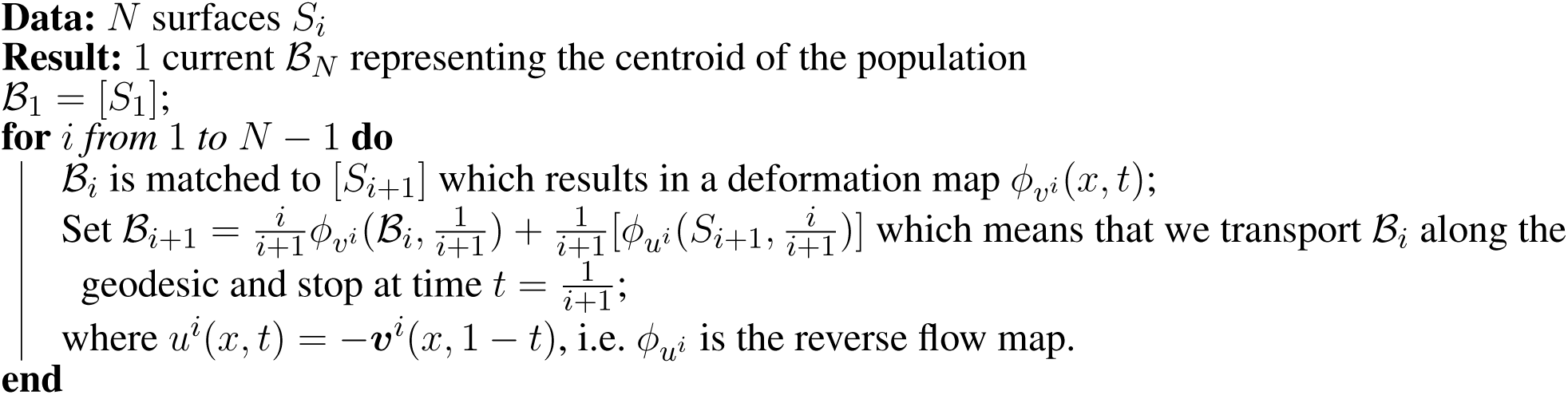

The weights in the averaging reflect the relative importance of the new shape, so that at the end of the procedure, all shapes forming the centroid have equal weight 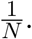

Note that we have used the notation 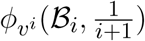 to denote the transport (push-forward) of the current 𝓑_*i*_ by the diffeomorphism. Here 𝓑_*i*_ is a linear combination of currents associated to surfaces, and the transported current is the linear combination (keeping the weights unchanged) of the currents associated to the transported surfaces.

### 3.4 Alternative method: Pairwise Centroid (PW)

Another possibility is to recursively split the population in two parts until having only one surface in each group (see Fig. 1), and then going back up along the dyadic tree by computing pairwise centroids between groups, with appropriate weight for each centroid (Algorithm 3).

**Figure 1.**
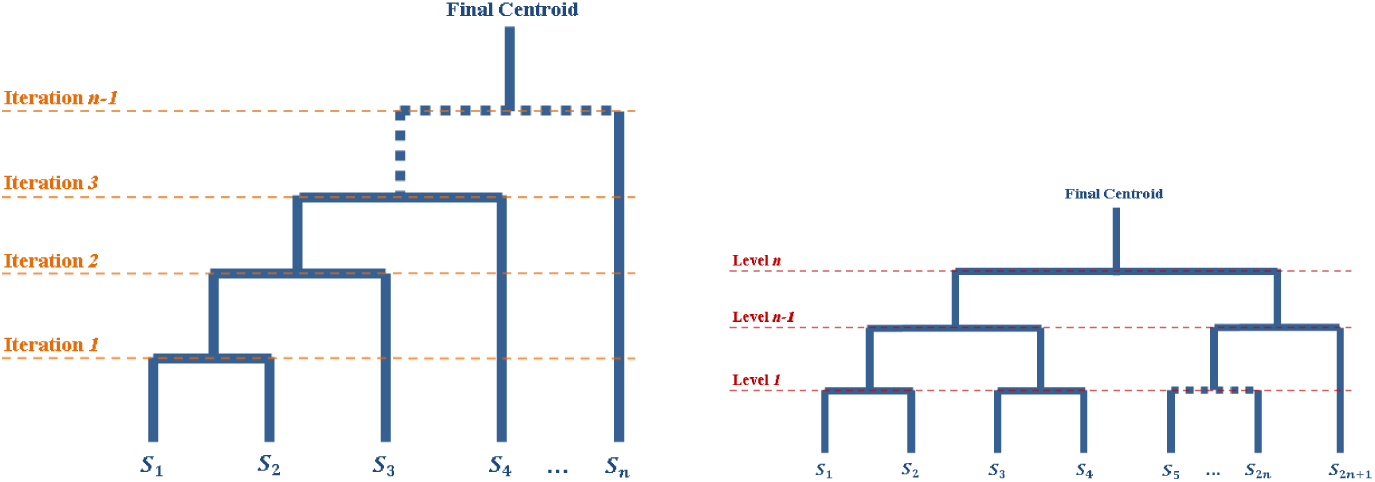
Diagrams of the iterative processes which lead to the centroids computations. The tops of the diagrams represent the final centroid. The diagram on the left corresponds to the Iterative Centroid algorithms (IC1 and IC2). The diagram on the right corresponds to the pairwise algorithm (PW).

#### Algorithm 3: Pairwise Centroid (PW)

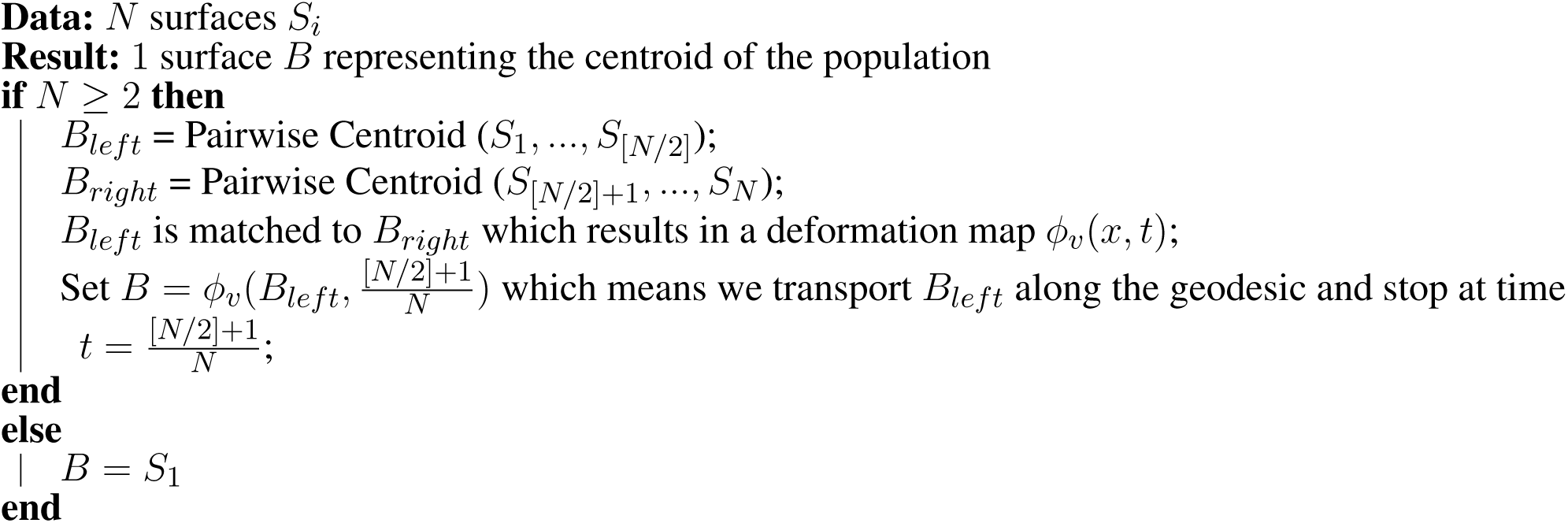

These three methods depend on the ordering of subjects. In a previous work (12), we showed empirically that different orderings result in very similar final centroids. Here we focus on the use of such centroid for statistical shape analysis.

### 3.5 Comparison with a variational template estimation method

In this study, we will compare our centroid approaches to a variational template estimation method proposed by Glaunés et al (10). This variational method estimates a template given a collection of surfaces using the framework of currents. It is posed as a minimum mean squared error estimation problem. Let *S_i_* be *N* surfaces in ℝ^3^ (i.e. the whole surface population). Let [*S_i_*] be the corresponding current of *S_i_*, or its approximation by a finite sum of vectorial Diracs. The problem is formulated as follows:

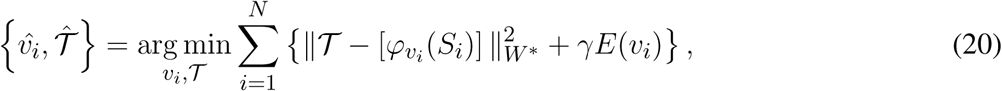

The method uses an alternated optimization i.e. surfaces are successively matched to the template, then the template is updated and this sequence is iterated until convergence. One can observe that when *φ_i_* is fixed, the functional is minimized when 𝓣 is the average of 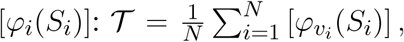 which makes the optimization with respect to 𝓣 straightforward. This optimal current is the union of all surfaces *φ_v_i__* (*S_i_*). However, all surfaces being co-registered, the *φ̂_v_i__*(*S_i_*) are close to each other, which makes the optimal template 𝓣̂ close to being a true surface. Standard initialization consists in setting 𝓣 = 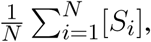 which means that the initial template is defined as the combination of all unregistered shapes in the population. Alternatively, if one is given a good initial guess 𝓣, the convergence speed of the method can be improved. In particular, the initialisation can be provided by iterative centroids; this is what we will use in the experimental section.

Regarding the computational complexity, the different centroid approaches perform *N* − 1 matchings while the variational template estimation requires *N* × *iter* matchings, where *iter* is the number of iterations. Moreover the time for a given matching depends quadratically on the number of vertices of the surfaces being matched. It is thus more expensive when the template is a collection of surfaces as in IC2 and in the variational template estimation.

## 4 Statistical Analysis

The proposed iterative centroid approaches can be used for subsequent statistical shape analysis of the population, using various strategies. A first strategy consists in analysing the deformations between the centroid and the individual subjects. This is done by analysing the initial momentum vectors 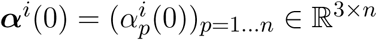 which encode the optimal diffeomorphisms computed from the matching between a centroid and the subjects *S_i_*. Initial momentum vectors all belong to the same vector space and are located on the vertices of the centroid. Different approaches can be used to analyse these momentum vectors, including Principal Component Analysis for the description of populations, Support Vector Machines or Linear Discriminant Analysis for automatic classification of subjects. A second strategy consists in analysing the set of pairwise distances between subjects. Then, the distance matrix can be entered into analysis methods such as Isomap (29), Locally Linear Embedding (30), (31) or spectral clustering algorithms (32). Here, we tested two approaches: i) the analysis of initial momentum vectors using a Kernel Principal Component Analysis for the first strategy; ii) the approximation of pairwise distance matrices for the second strategy. These tests allow us both to validate the different iterative centroid methods and to show the feasibility of such analysis on large databases.

### 4.1 Principal Component Analysis on initial momentum vectors

The Principal Component Analysis (PCA) on initial momentum vectors from the template to the subjects of the population is an adaptation of PCA in which Euclidean scalar products between observations are replaced by scalar products using a kernel. Here the kernel is *K_V_* the kernel of the R.K.H.S *V*. This adaptation can be seen as a Kernel PCA (13). PCA on initial momentum vectors has previously been used in morphometric studies in the LDDMM setting (3, 33) and it is sometimes referred to tangent PCA.

We briefly recall that, in standard PCA, the principal components of a dataset of *N* observations ***a**^i^* ∈ ℝ^*P*^ with *i* ∈ {1, …, *N*} are defined by the eigenvectors of the covariance matrix *C* with entries:

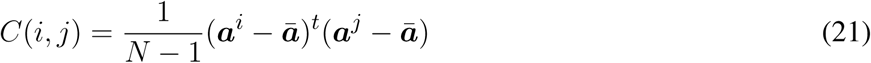

with *a^i^* given as a column vector, 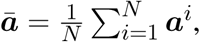 and *x^t^* denotes the transposition of a vector ***x***.

In our case, our observations are initial momentum vectors ***α**^i^* ∈ ℝ^3×*n*^ and instead of computing the Euclidean scalar product in ℝ^3×*n*^, we compute the scalar product with matrix *K_V_*, which is a natural choice since it corresponds to the inner product of the corresponding initial vector fields in the space *V*. The covariance matrix then writes:

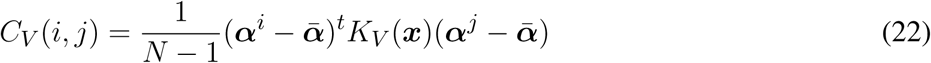

with ***ᾱ*** the vector of the mean of momentum vectors, and ***x*** the vector of vertices of the template surface. We denote λ_1_, λ_2_, …, λ_*N*_ the eigenvalues of *C* in decreasing order, and ***ν***^1^, ***ν***^2^, …, ***ν**^N^* the corresponding eigenvectors. The *k*-*th* principal mode is computed from the *k*-*th* eigenvector *ν^k^* of *C_V_*, as follows:

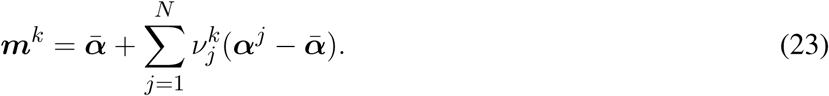

The cumulative explained variance *CEV_k_* for the *k* first principal modes is given by the equation:

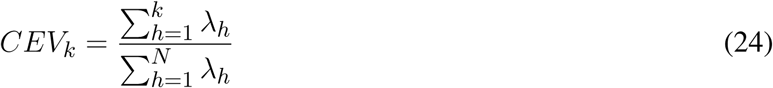

We can use geodesic shooting along any principal mode ***m**^k^* to visualise the corresponding deformations.

#### Remark

To analyse the population, we need to know the initial momentum vectors ***α**^i^* which correspond to the matchings from the centroid to the subjects. For the IC1 and PW centroids, these initial momentum vectors were obtained by matching the centroid to each subject. For the IC2 centroid, since the mesh structure is composed of all vertices of the population, it is too computationally expensive to match the centroid toward each subject. Instead, from the deformation of each subject toward the centroid, we used the opposite vector of final momentum vectors for the analysis. Indeed, if we have two surfaces *S* and *T* and need to compute the initial momentum vectors from *T* to *S*, we can estimate the initial momentum vectors ***α**^TS^*(0) from *T* to *S* by computing the deformation from *S* to *T* and using the initial momentum vectors ***α**̃^TS^*(0) = −***α**^ST^*(1), which are located at vertices *ϕ_ST_*(*x^S^*).

### 4.2 Distance matrix approximation

Various methods such as Isomap (29) or Locally Linear Embedding (30) (31) use as input a matrix of pairwise distances between subjects. In the LDDMM setting, it can be computed using diffeomorphic distances: *ρ*(*S_i_*, *S_j_*) = *D*(*id*, *φ_ij_*). However, for large datasets, computing all pairwise deformation distance is computationally very expensive, as it involves *O*(*N*^2^) matchings. An alternative is to approximate the pairwise distance between two subjects through their matching from the centroid or template. This approach has been introduced in Yang et al (34). Here we use a first order approximation to estimate the diffeomorphic distance between two subjects:

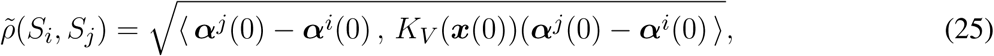

with ***x***(0) the vertices of the estimated centroid or template and ***α**^i^* (0) is the vector of initial momentum vectors computed by matching the template to *S_i_*. Using such approximation allows to compute only *N* matchings instead of *N*(*N* − 1).

Note that *ρ*(*S_i_*, *S_j_*) is in fact the distance between *S_i_* and *φ_ij_*(*S_i_*), and not between *S_i_* and *S_j_* due to the not exactitude of matchings. However we will refer to it as a distance in the following to denote the dissimilarity between *S_i_* and *S_j_*.

## 5 Experiments and Results

In this section, we evaluate the use of iterative centroids for statistical shape analysis. Specifically, we investigate the centring of the centroids within the population, their impact on population analysis based on Kernel PCA and on the computation of distance matrices. For our experiments, we used three different datasets: two real datasets and a synthetic one. In all datasets shapes are hippocampi. The hippocampus is an anatomical structure of the temporal lobe of the brain, involved in different memory processes.

### 5.1 Data

The two real datasets are from the European database IMAGEN (36) ^1^ composed of young healthy subjects. We segmented the hippocampi from T1-weighted Magnetic Resonance Images (MRI) of subjects using the SACHA software (35) (see Fig. 2). The synthetic dataset was built using deformations of a single hippocampal shape of the IMAGEN database.

**Figure 2.**
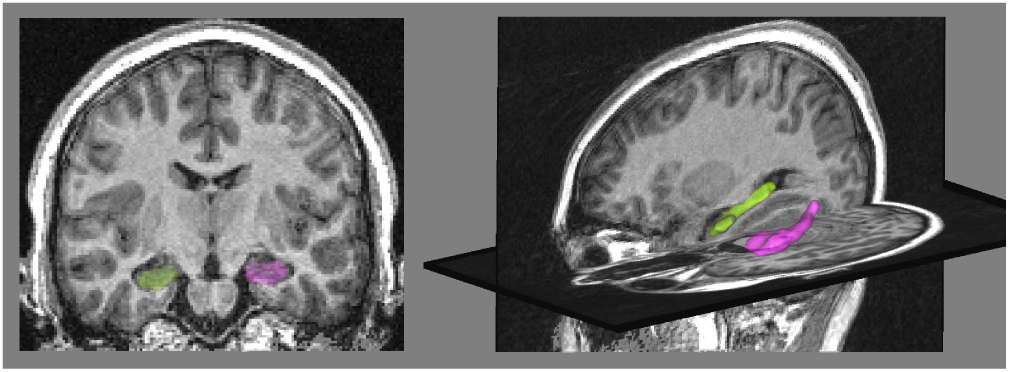
Left panel: coronal view of the MRI with the binary masks of hippocampi segmented by the SACHA software (35), the right hippocampus is in green and the left one in pink. Right panel: 3D view of the hippocampus meshes.

### The synthetic dataset SD

is composed of synthetic deformations of a single shape *S*_0_, designed such that this single shape becomes the exact center of the population. We will thus be able to compare the computed centroids to this exact center. We generated 50 subjects for this synthetic dataset from *S*_0_, along geodesics in different directions. We randomly chose two orthogonal momentum vectors *β*_1_ and *β*_2_ in ℝ^3×*n*^. We then computed momentum vectors ***α**^i^*, *i* ∈ {1, …, 25} of the form 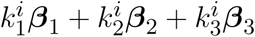 with 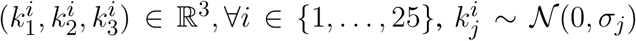 with *σ*_1_ > *σ*_2_ ≫ *σ*_3_ and *β*_3_ a randomly selected momentum vector, adding some noise to the generated 2D space. We computed momentum vectors ***α**^j^*, *j* ∈ {26, …, 50} such as ***α**^j^* = −***α***^*j*−25^. We generated the 50 subjects of the population by computing geodesic shootings of *S*_0_ using the initial momentum vectors ***α**^i^*, *i* ∈ {1, …, 50}. The population is symmetrical since 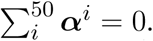 It should be noted that all shapes of the dataset have the same mesh structure composed of *n* = 549 vertices.

#### The real dataset RD50

is composed of 50 left hippocampi from the IMAGEN database. We applied the following preprocessing steps to each individual MRI. First, the MRI was linearly registered toward the MNI152 atlas, using the FLIRT procedure (37) of the FSL software ^2^. The computed linear transformation was then applied to the binary mask of the hippocampal segmentation. A mesh of this segmentation was then computed from the binary mask using the BrainVISA software ^3^. All meshes were then aligned using rigid transformations to one subject of the population. For this rigid registration, we used a similarity term based on measures (as in (38)). All meshes were decimated in order to keep a reasonable number of vertices: meshes have on average 500 vertices.

#### The real database RD1000

is composed of 1000 left hippocampi from the IMAGEN database. We applied the same preprocessing steps to the MRI data as for the dataset RD50. This dataset has also a score of Incomplete Hippocampal Inversion (IHI) (14), which is an anatomical variant of the hippocampus, present in 17% of the normal population.

### 5.2 Experiments

For the datasets SD and RD50 (which both contain 50 subjects), we compared the results of the three different iterative centroid algorithms (IC1, IC2 and PW). We also investigated the possibility of computing variational templates, initialized by the centroids, based on the approach presented in section 3.5. We could thus compare the results obtained when using the centroid directly to those obtained when using the most expensive (in term of computation time) template estimation. We thus computed 6 different centres: IC1, IC2, PW and the corresponding variational templates T(IC1), T(IC2), T(PW). For the synthetic dataset SD, we could also compare those 6 estimated centres to the exact centre of the population. For the real dataset RD1000 (with 1000 subjects), we only computed the iterative centroid IC1.

For all computed centres and all datasets, we investigated: 1) the computation time; 2) whether the centres are close to a critical point of the Fréchet functional of the manifold discretised by the population; 3) the impact of the estimated centres on the results of Kernel PCA; 4) their impacts on approximated distance matrices.

To assess the “centring” (i.e. how close an estimated centre is to a critical point of the Fréchet functional) of the different centroids and variational templates, we computed a ratio using the momentum vectors from centres to subjects. The ratio *R* takes values between 0 and 1:

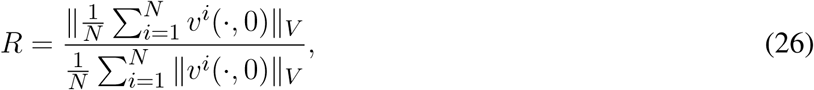

with *v^i^*(⋅; 0) the vector field of the deformation from the estimated centre to the subject *S_i_*, corresponding to the vector of initial momentum vectors ***α**^i^*(0).

We compared the results of Kernel PCA computed from these different centres by comparing the principal modes and the cumulative explained variance for different number of dimensions.

Finally, we compared the approximated distance matrices to the direct distance matrix.

For the RD1000 dataset, we will try to predict an anatomical variant of the normal population, the Incomplete Hippocampal Inversion (IHI), presents in only 17% of the population.

### 5.3 Synthetic dataset SD

#### 5.3.1 Computation time

All the centroids and variational templates have been computed with *σ_V_* = 15, which represents roughly half of the shapes length. Computation times for IC1 took 31 minutes, 85 minutes for IC2, and 32 minutes for PW. The corresponding variational template initialised by these estimated centroids took 81 minutes (112 minutes in total), 87 minutes (172 minutes in total) and 81 minutes (113 minutes in total). As a reference, we also computed a template with the standard initialisation whose computation took **194** minutes. Computing a centroid saved between 56% and 84% of computation time over the template with standard initialization and between 50% and 72% over the template initialized by the centroid.

#### 5.3.2 “Centring” of the estimated centres

Since in practice a computed centre is never at the exact centre, and its estimation may vary accordingly to the discretisation of the underlying shape space, we decided to generate another 49 populations, so we have 50 different discretisations of the shape space. For each of these populations, we computed the 3 centroids and the 3 variational templates initialized with these centroids. We calculated the ratio *R* described in the previous section for each estimated centre. Table 1 presents the mean and standard deviation values of the ratio *R* for each centroid and template, computed over these 50 populations.

**Table 1.** Synthetic dataset SD. Ratio *R* (equation 26) computed for the 3 centroids *C*, and the 3 variational templates initialized via these centroids (*T*(*C*)).

| Ratio | $C$ | $T(C)$ |
| --- | --- | --- |
| IC1 | $0.07 \pm 0.03$ | $0.05 \pm 0.02$ |
| IC2 | $0.07 \pm 0.03$ | $0.05 \pm 0.02$ |
| PW | $0.11 \pm 0.05$ | $0.07 \pm 0.02$ |

In a pure Riemannian setting (i.e. disregarding the fact that matchings are not exact), a zero ratio would mean that we are at a critical point of the Fréchet functional, and under some reasonable assumptions on the curvature of the shape space in the neighbourhood of the dataset (which we cannot check however), it would mean that we are at the Fréchet mean. By construction, the ratio computed from the exact centre using the initial momentum vectors ***α**^i^* used for the construction of subjects *S_i_* (as presented in section 5.1) is zero.

Ratios *R* are close to zero for all centroids and variational templates, indicating that they are close to the exact centre. Furthermore, the value of *R* may be partly due to the non-exactitude of the matchings between the estimated centres and the subjects. To become aware of this non-exactitude, we matched the exact centre toward all subjects of the SD dataset. The resulting ratio is *R* = 0.05. This is of the same order of magnitude as the ratios obtained in Table 1, indicating that the estimated centres are indeed very close to the exact centre.

#### 5.3.3 PCA on initial momentum vectors

We performed a PCA computed with the initial momentum vectors (see section 4 for details) from our different estimated centres (3 centroids, 3 variational templates and the exact centre).

We computed the cumulative explained variance for different number of dimensions of the PCA. Results are presented in Table 2. The cumulative explained variances are very similar for the different centres for any number of dimensions.

**Table 2.** Synthetic dataset SD. Proportion of cumulative explained variance of kernel PCA computed from different centres.

|  | 1st mode | 2nd mode | 3rd mode |
| --- | --- | --- | --- |
| Centre | 0.829 | 0.989 | 0.994 |
| IC1 | 0.829 | 0.990 | 0.995 |
| IC2 | 0.833 | 0.994 | 0.996 |
| PW | 0.829 | 0.990 | 0.995 |
| T(IC1) | 0.829 | 0.995 | 0.999 |
| T(IC2) | 0.829 | 0.995 | 0.999 |
| T(PW) | 0.829 | 0.995 | 0.999 |

We wanted to take advantage of the construction of this synthethic dataset to answer the question: Do the principal components explain the same deformations? The SD dataset allows to visualise the principal component relatively to the real position of the generator vectors and the population itself. Such visualisation is not possible for real dataset since shape spaces are not generated by only 2 vectors. For this synthetic dataset, we can project principal components on the 2D space spanned by *β*_1_ and *β*_2_ as described in the previous paragraph. This projection allows displaying in the same 2D space subjects in their native space, and principal axes computed from the different Kernel PCAs. To visualize the first component (respectively the second one), we shot from the associated centre in the direction *k* * ***m***^1^ (resp. ***m***^2^) with 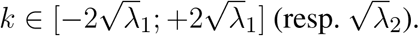 Results are presented in Figure 3. The deformations captured by the 2 principal axes are extremely similar for all centres. The principal axes for the 7 centres, have all the same position within the 2D shape space. So for a similar amount of explained variance, the axes describe the same deformation.

**Figure 3.**
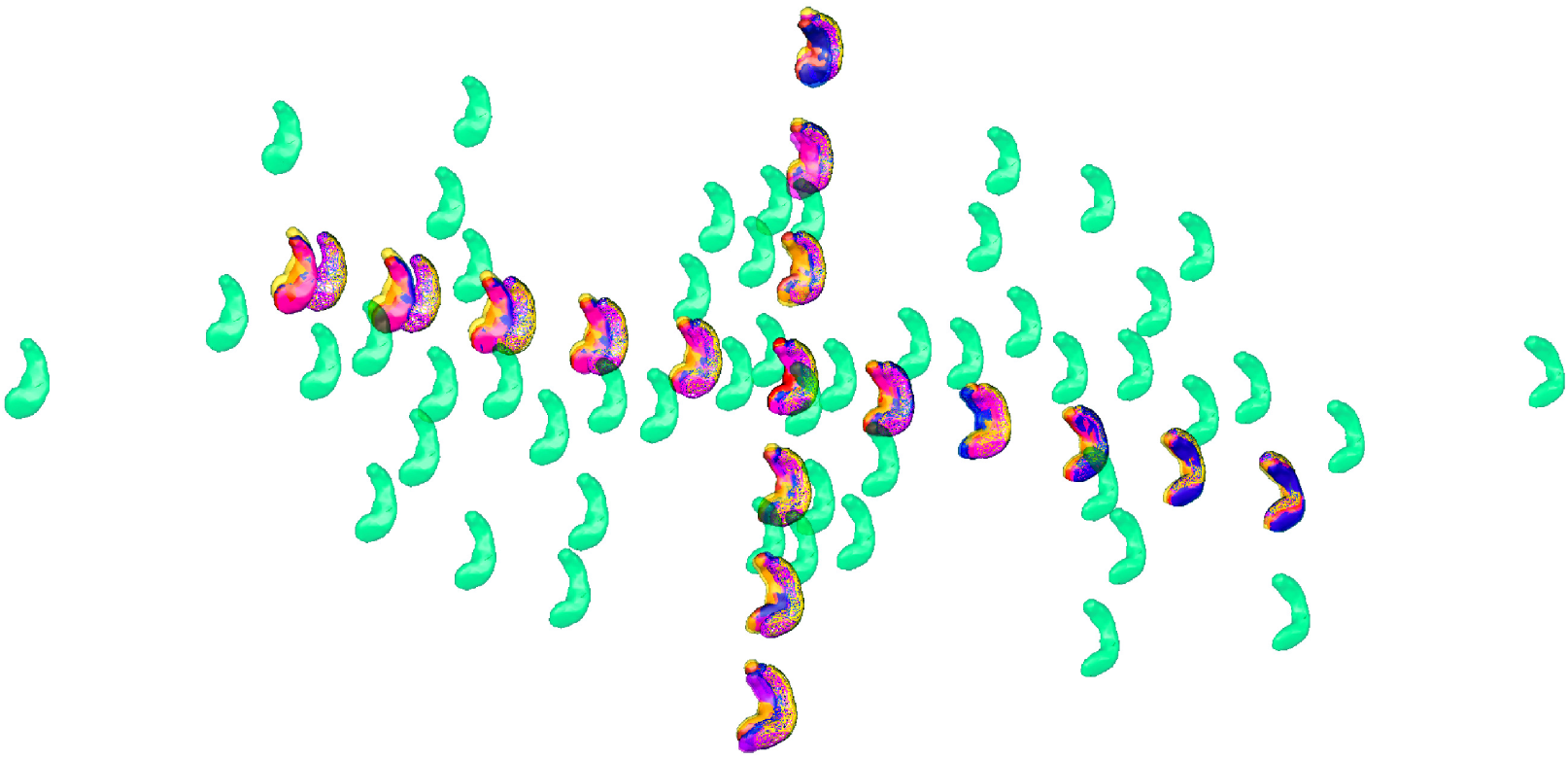
Synthetic dataset SD. Illustration of the two principal components of the 6 centres projected into the 2D space of the SD dataset. Synthetic population in green, the real centre is in red in the middle. The two axes are the two principal components projected into the 2D space of the population computed from 7 different estimated centres (marked in orange for the exact centre, in blue for IC1, in yellow for IC2 and in magenta for PW), they go from −2*σ* to +2*σ* of the variability of the corresponding axe. This figure shows that the 6 different tangent spaces projected into the 2D space of shapes, are very similar, even if the centres have different positions.

Overall, for this synthetic dataset, the 6 estimated centres give very similar PCA results.

#### 5.3.4 Distance matrices

We then studied the impact of different centres on the approximated distance matrices. We computed the seven approximated distance matrices corresponding to the seven centres, and the direct pairwise distance matrix computed by matching all subjects to each other. Computation of the direct distance matrix took 1000 minutes (17 hours) for this synthetic dataset of 50 subjects. In the following, we denote as *aM*(*C*) the approximated distance matrix computed from the centre *C*.

To quantify the difference between these matrices, we used the following error *e*:

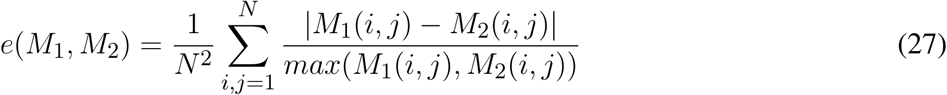

with *M*_1_ and *M*_2_ two distance matrices. Results are reported in Table 3. For visualisation of the errors against the pairwise computed distance matrices, we also computed the error between the direct distance matrix, by computing pairwise deformations (23h hours of computation per population), for 3 populations randomly selected. Figure 4 shows scattered plots between the pairwise distances matrices and the approximated distances matrices of IC1 and T(IC1) for the 3 randomly selected populations. The errors between the *aM*(*IC*1) and the pairwise distance matrices of each of the populations are 0.17 0.16 and 0.14, respectively 0.11 0.08 and 0.07 for the errors with the corresponding *aM*(*T*(*IC*1)). We can observe a subtle curvature of the scatter-plot, which is due to the curvature of the shape space. This figure illustrates the good approximation of the distances matrices, regarding to the pairwise estimation distance matrix. The variational templates are getting slightly closer to the identity line, which is expected (as for the better ratio values) since they have extra iterations to converge to a centre of the population, however the estimated centroids from the different algorithms, still provide a good approximation of the pairwise distances of the population. In conclusion for this set of synthetic population, the different estimated centres have also a little impact on the approximation of the distance matrices.

**Figure 4.**
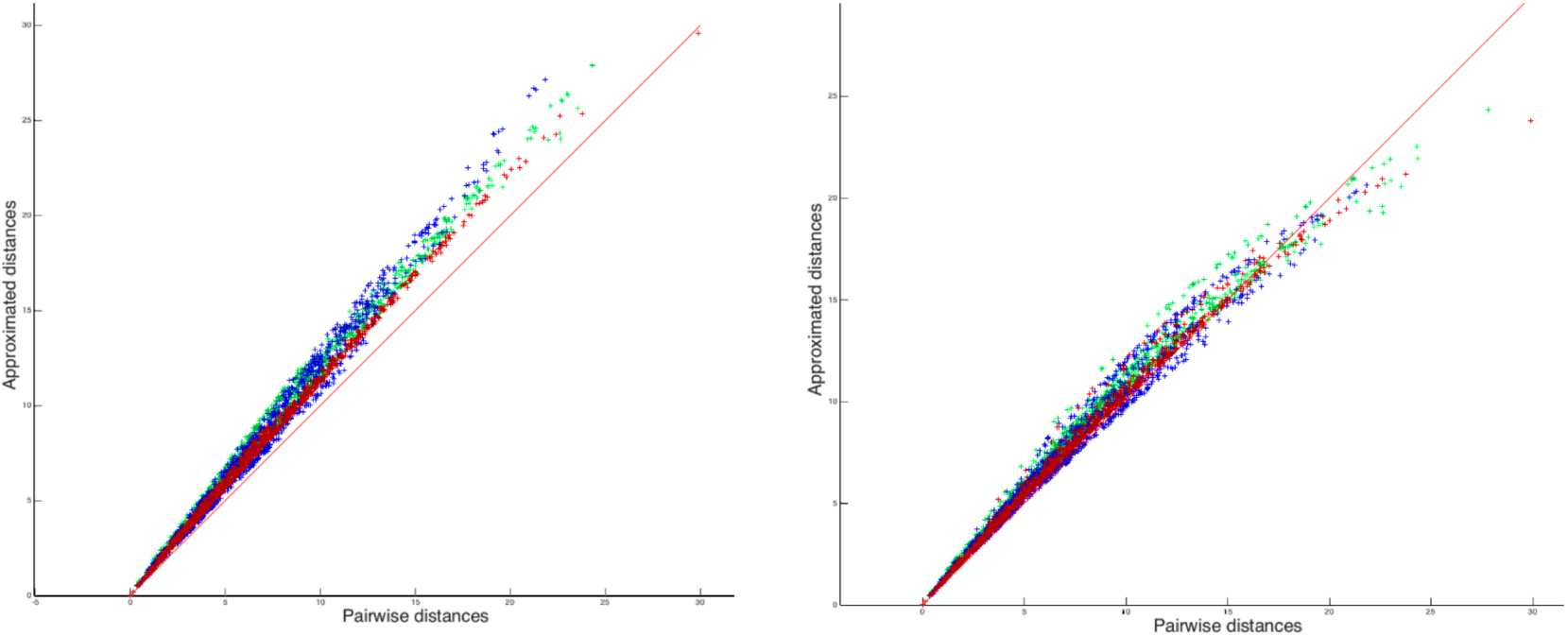
Synthetic datasets generated from SD50. Left: scatter plot between the approximated distance matrices *aM*(*T*(*IC*1)) of 3 different populations, and the pairwise distances matrices of the corresponding population. Right: scatter plot between the approximated distance matrices *aM*(*IC*1) of 3 different populations, and the pairwise distances matrices of the corresponding population. The red line corresponds to the identity.

**Table 3.** Synthetic dataset SD. Mean ± standard deviation errors *e* (equation 27) between the six different approximated distance matrices for each of the generated data sets.

| $e(aM(\cdot), aM(\cdot))$ | IC2 | PW | T(IC1) | T(IC2) | T(PW) |
| --- | --- | --- | --- | --- | --- |
| IC1 | $0.001 \pm 0.001$ | $0.002 \pm 0.002$ | $0.084 \pm 0.02$ | $0.084 \pm 0.02$ | $0.084 \pm 0.02$ |
| IC2 | 0 | $0.003 \pm 0.002$ | $0.084 \pm 0.02$ | $0.084 \pm 0.02$ | $0.084 \pm 0.02$ |
| PW | | 0 | $0.084 \pm 0.02$ | $0.084 \pm 0.02$ | $0.084 \pm 0.02$ |
| T(IC1) | | | 0 | $0.001 \pm 0.001$ | $0.002 \pm 0.001$ |
| T(IC2) | | | | 0 | $0.002 \pm 0.001$ |

### 5.4 The real dataset RD50

We now present experiments on the real dataset RD50. For this dataset, the exact center of the population is not known, neither is the distribution of the population and meshes have different numbers of vertices and different connectivity structures.

#### 5.4.1 Computation time

We estimated our 3 centroids IC1 (75 minutes) IC2 (174 minutes) and PW (88 minutes), and the corresponding variational templates, which took respectively 188 minutes, 252 minutes and 183 minutes. The total computation time for *T*(*IC*1) is 263 minutes, 426 minutes for *T*(*IC*2) and 271 minutes for *T*(*PW*).

For comparison of computation time, we also computed a template using the standard initialization (the whole population as initialisation) which took 1220 minutes (20.3 hours). Computing a centroid saved between 85% and 93% of computation time over the template with standard initialization and between 59% and 71% over the template initialized by the centroid.

#### 5.4.2 Centring of the centres

As for the synthetic dataset, we assessed the centring of these six different centres. To that purpose, we first computed the ratio *R* of equation (26), for the centres estimated via the centroids methods, IC1 has a *R* = 0.25, for IC2 the ratio is *R* = 0.33 and for PW it is *R* = 0.32. For centres estimated via the variational templates initialised by those centroids, the ratio for T(IC1) is *R* = 0.21, for T(IC2) is *R* = 0.31 and for T(PW) is *R* = 0.26.

The ratios are higher than for the synthetic dataset indicating that centres are less centred. This was predictable since the population is not built from one surface via geodesic shootings as the synthetic dataset. In order to better understand these values, we computed the ratio for each subject of the population (after matching each subject toward the population), as if each subject was considered as a potential centre. For the whole population, the average ratio was 0.6745, with a minimum of 0.5543, and a maximum of 0.7626. These ratios are larger than the one computed for the estimated centres and thus the 6 estimated centres are closer to a critical point of the Fréchet functional than any subject of the population.

#### 5.4.3 PCA on initial momentum vectors

As for the synthetic dataset, we performed six PCAs from the estimated centres.

Figure 5 and Table 4 show the proportion of cumulative explained variance for different number of modes. We can note that for any given number of modes, all PCAs result in the same proportion of explained variance.

**Figure 5.**
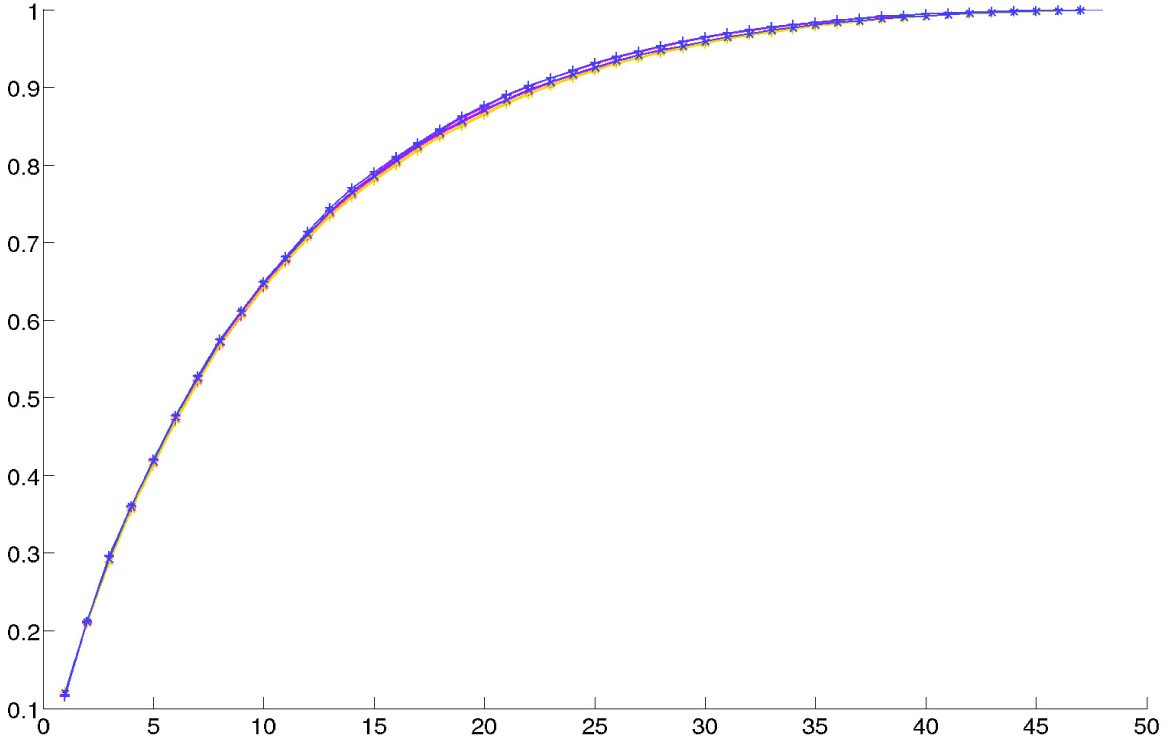
Real dataset RD50. Proportion of cumulative explained variance for Kernel PCAs computed from the 6 different centres, with respect to the number of dimensions. Curves are almost identical.

**Table 4.** Real dataset RD50. Proportion of cumulative explained variance for kernel PCAs computed from the 6 different centres, for different number of dimensions

|  | 1st mode | 2nd mode | 15th mode | 20th mode |
| --- | --- | --- | --- | --- |
| IC1 | 0.118 | 0.214 | 0.793 | 0.879 |
| IC2 | 0.121 | 0.209 | 0.780 | 0.865 |
| PW | 0.117 | 0.209 | 0.788 | 0.875 |
| T(IC1) | 0.117 | 0.222 | 0.815 | 0.899 |
| T(IC2) | 0.115 | 0.220 | 0.814 | 0.898 |
| T(PW) | 0.116 | 0.221 | 0.814 | 0.898 |

#### 5.4.4 Distance matrices

As for the synthetic dataset, we then studied the impact of these different centres on the approximated distance matrices. A direct distance matrix was also computed (around 90 hours of computation time). We compared the approximated distance matrices of the different centres to: i) the approximated matrix computed with IC1; ii) the direct distance matrix.

We computed the errors *e*(*M*_1_, *M*_2_) defined in equation 27. Results are presented in Table 5. Errors are small and with the same order of magnitude.

**Table 5.**
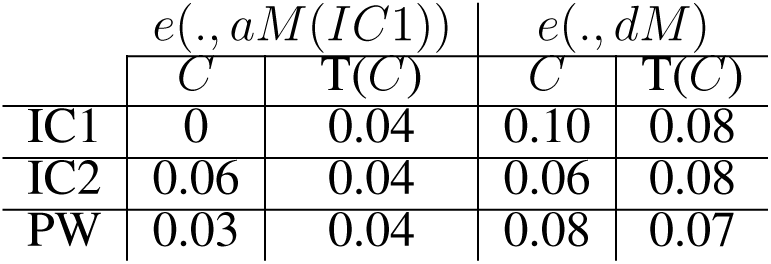
Real dataset RD50. Errors *e* (equation 27) between the approximated distance matrices of each estimated centre and: i) the approximated matrix computed with IC1 (left columns); ii) the direct distance matrix (right columns).

| | $e(\cdot, aM(IC1))$ | | $e(\cdot, dM)$ | |
| --- | --- | --- | --- | --- |
| | $C$ | $T(C)$ | $C$ | $T(C)$ |
| IC1 | 0 | 0.04 | 0.10 | 0.08 |
| IC2 | 0.06 | 0.04 | 0.06 | 0.08 |
| PW | 0.03 | 0.04 | 0.08 | 0.07 |

Figure 6a shows scatter plots between the direct distance matrix and the six approximated distance matrices. Interestingly, we can note that the results are similar to those obtained by Yang et al. ((34), Figure 2). Figure 6b shows scatter plots between the approximated distance matrix from IC1 and the five others approximated distance matrices. The approximated matrices thus seem to be largely independent of the chosen centre.

**Figure 6.**
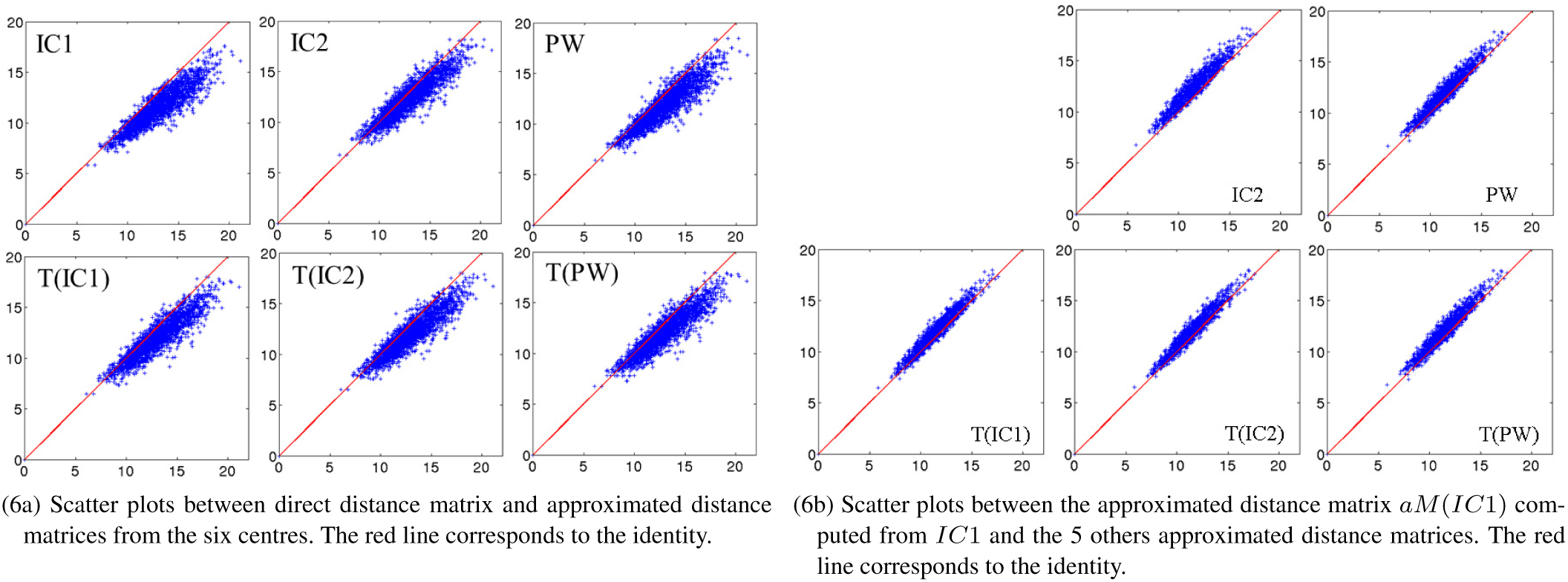
Real dataset RD50. Scatter plots

### 5.5 Real dataset RD1000

Results on the real dataset RD50 and the synthetic SD showed that results were highly similar for the 6 different centres. In light of these results and because of the large size of the real dataset RD1000, we only computed IC1 for this last dataset. The computation time was about 832 min (13.8 hours) for the computation of the centroid using the algorithm IC1, and 12.6 hours for matching the centroid to the population.

The ratio *R* of equation 26 computed from the IC1 centroid was 0.1011, indicating that the centroid is well centred within the population.

We then performed a Kernel PCA on the initail momentum vectors from this IC1 centroid to the 1000 shapes of the population. The proportions of cumulative explained variance from this centroid are 0.07 for the 1st mode, 0.12 for the 2nd mode, 0.48 for the 10th mode, 0.71 for the 20th mode, 0.85 for the 30th mode, 0.93 for the 40th mode, 0.97 for the 50th mode and 1.0 from the 100th mode. In addition, we explored the evolution of the cumulative explained variance when considering varying numbers of subjects in the analysis. Results are displayed in Figure 7. We can first note that about 50 dimensions are sufficient to describe the variability of our population of hippocampal shapes from healthy young subjects. Moreover, for large number of subjects, this dimensionality seems to be stable. When considering increasing number of subjects in the analysis, the dimension increases and converges around 50.

**Figure 7.**
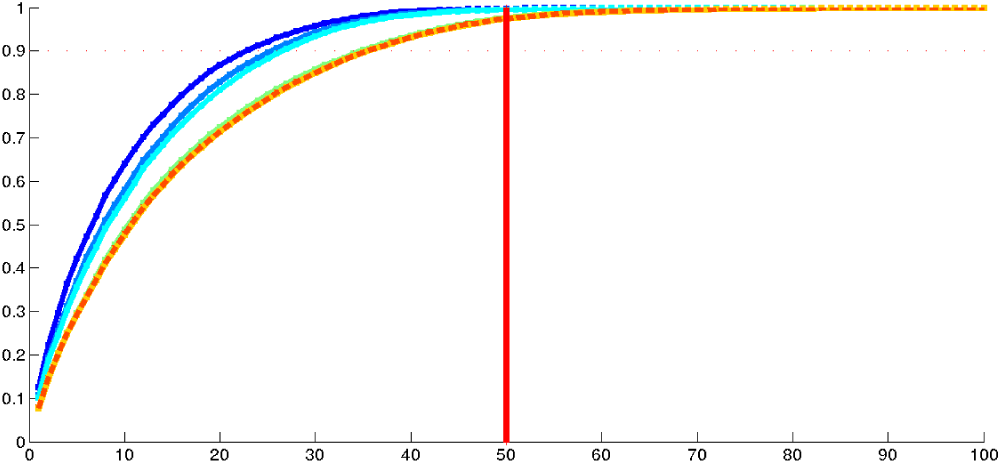
Real dataset RD1000. Proportion of cumulative explained variance of K-PCA as a function of the number of dimensions (in abscissa) and considering varying number of subjects. The dark blue curve was made using 100 subjects, the blue 200, the light blue 300, the green curve 500 subjects, the yellow one 800, very close to the dotted orange one which was made using 1000 subjects.

Finally, we computed the approximated distance matrix. Its histogram is shown in Figure 8. It can be interesting to note that, as for RD50, the average pairwise distance between the subject is around 12, which means nothing by itself, but the points cloud on Figure 6a and the histogram on Figure 8, show no pairwise distances below 6, while the minimal pairwise distance for the SD50 dataset - generated by a 2D space - is zero. This corresponds to the intuition that, in a space of high dimension, all subjects are relatively far from each other.

**Figure 8.**
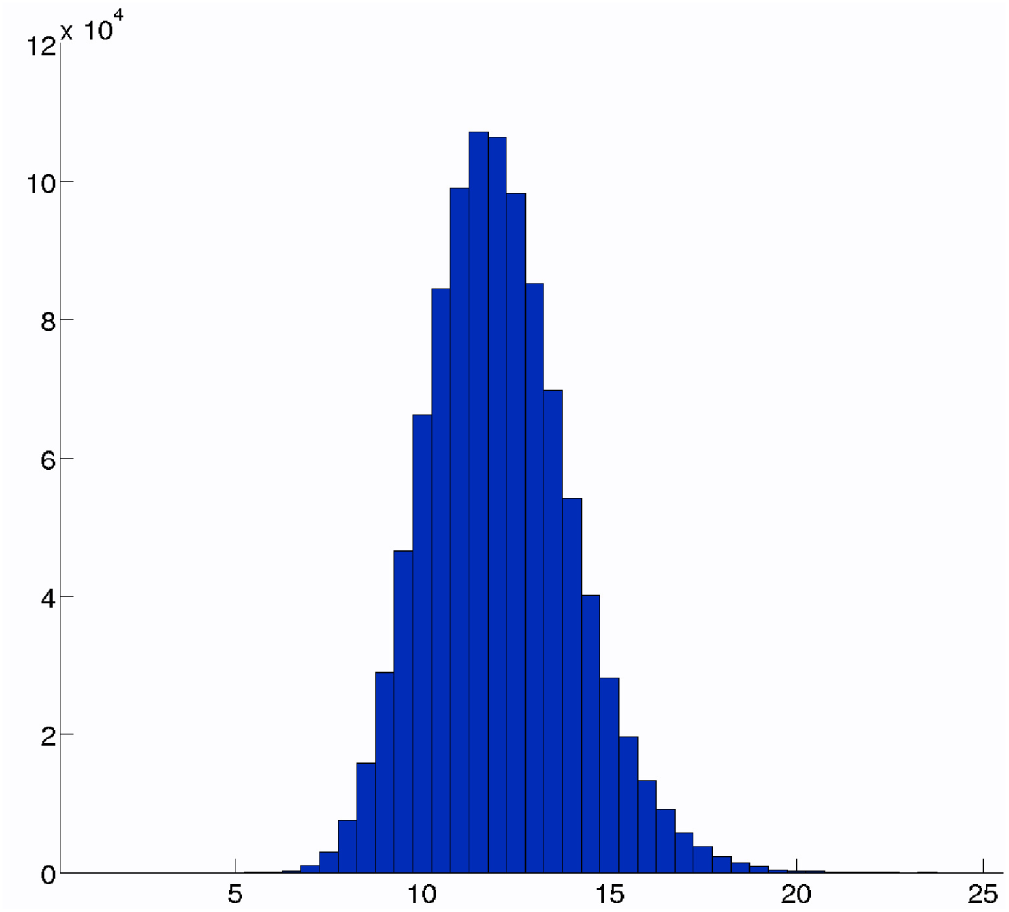
Real dataset RD1000. Histogram of the approximated distances of the large database from the computed centroid.

#### 5.5.1 Prediction of IHI using shape analysis

Incomplete Hippocampal Inversion(IHI) is an anatomical variant of the hippocampal shape present in 17% of the normal population. All those 1000 subjects have an IHI score (ranking from 0 to 8) (14), which indicates how strong is the incomplete inversion of the hippocampus A null score means there is no IHI, 8 means there is a strong IHI. We now apply our approach to predict incomplete hippocampal inversions (IHI) from hippocampal shape parameters. Specifically, we predict the visual IHI score, which corresponds to the sum of the individual criteria as defined in (14). Only 17% of the population have an IHI score higher than 3.5. We studied whether it is possible to predict the IHI score using statistical shape analysis on the RD1000 dataset composed of 1000 healthy subjects (left hippocampus).

The deformation parameters characterising the shapes of the population are the eigenvectors computed from the centroid IC1, and they are the independent variables we will use to predict the IHI scores. As we saw in the previous step, 40 eigenvectors are enough to explain 93% of the total anatomical variability of the population. We use the centred and normalized principal eigenvectors *X*_1,*i*_, …, *X*_40,*i*_ computed from the RD1000 database with *i* ∈ {1, …, 1000} to predict the IHI score *Y*. We simply used a multiple linear regression model (39) which is written as 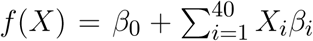 where *β*_0_,*β*_1_, … *β*_40_ are the regression coefficients to estimate. The standard method to estimate the regression coefficients is the least squares estimation method in which the coefficients *β_i_* minimize the residual sum of squares 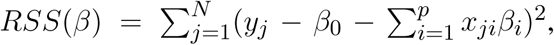 which leads to the estimated *β̂* (with matrix notations) *β̂* = (*X^T^X*)^−1^*X^T^Y*. For each number of dimensions *p* ∈ {1, …, 40} we validated the quality of the computed model with the adjusted coefficient of determination 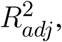 which expresses the part of explained variance of the model with respect to the total variance:

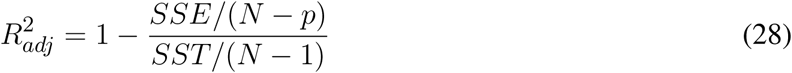

with 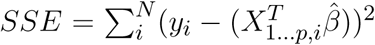 the residual variance due to the model and 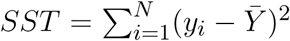 the total variance of the model. The 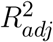 coefficient, unlike *R*^2^, takes into account the number of variables and therefore does not increase with the number of variables. One can note that *R* is the coefficient of correlation of Pearson. We then tested the significance of each model by computing the F statistic

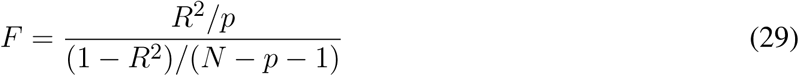

which follows a F-distribution with (*p*, *n* − *p* − 1) degrees of freedom. So for each number of variables (i.e. dimensions of the PCA space) we computed the adjusted coefficient of determination to evaluate the model and the *p*-value to evaluate the significance of the model.

Then we used the *k*-fold cross validation method which consists in using 1000 − *k* subjects to predict the *k* remaining ones. To quantify the prediction of the model, we used the traditional mean square error *MSE* = *SSE*/*N* which corresponds to the unexplained residual variance. For each model, we computed 10,000 *k*-fold cross validation and displayed the mean and the standard deviation of *MSE* corresponding to the model.

Results are given at Figure 9, and display the coefficient of determination of each model. The cross validation is only computed on models with a coefficient of correlation higher than 0.5, so models using at least 20 dimensions. For the *k*-fold cross validation, we chose *k* = 100 which represents 10% of the total population. Figure 9D presents results of cross validation; for each model computed from 20 to 40 dimensions we computed the mean of the 10,000 *MSE* of the 100-fold and its standard deviation. To have a point of comparison, we also computed the *MSE* between the IHI scores and random values which follow a normal distribution with the same mean and standard deviation as the IHI scores (red cross on the Figure). The *MSE* of the cross validation are similar to the *MSE* of the training set. This results show that using the first 30 to 40 principal components of initial momentum vectors computed from a centroid of the population, it is possible to predict the IHI score with a correlation of 69%. The firsts principal components (between 1 and 20) represent general variance maybe characteristic of the normal population, the shape differences related to IHI appear after. It is indeed expected that the principal (i.e. the first once) modes of variation does not capture a modification of the anatomy present in only 17% of the population.

**Figure 9.**
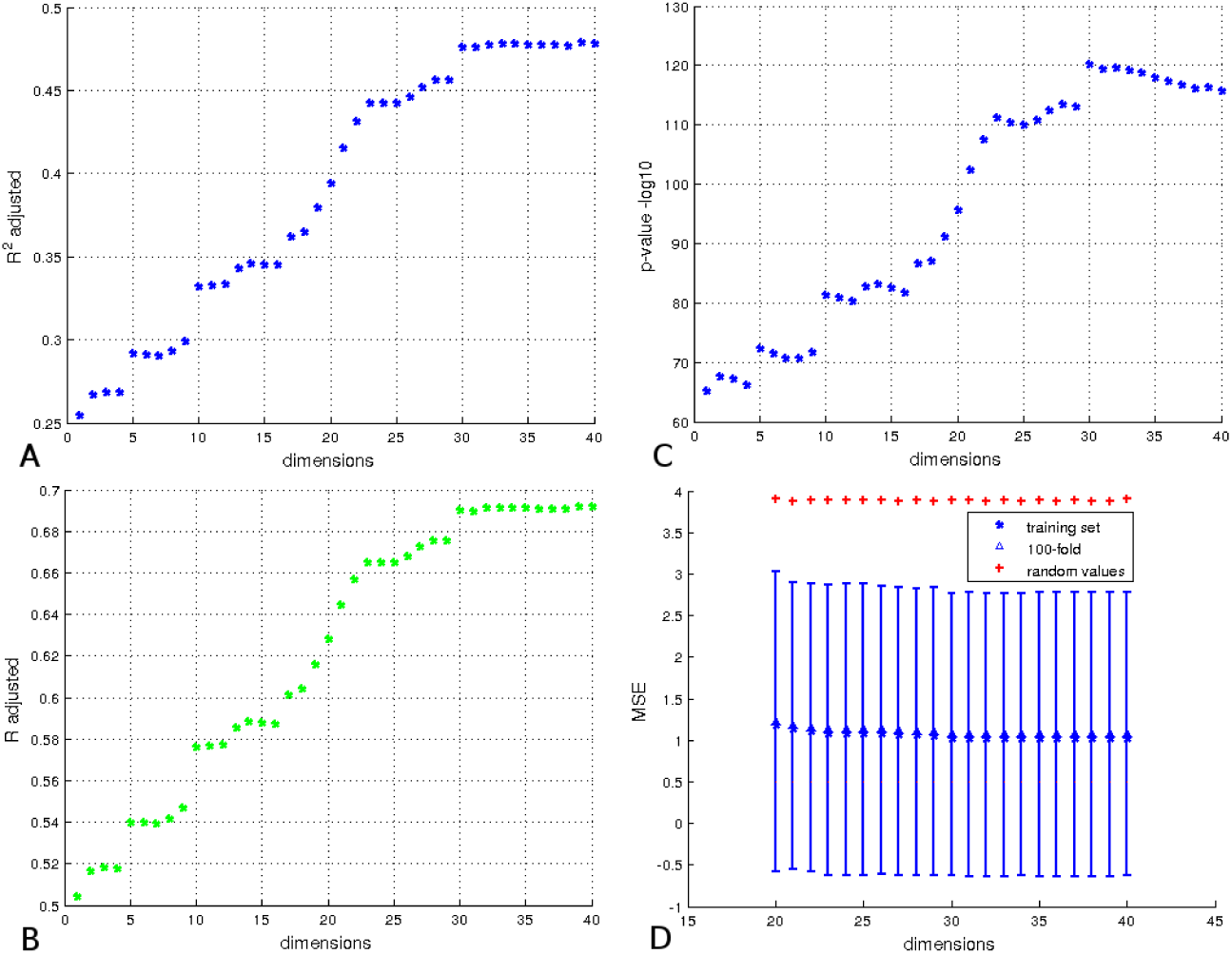
Results for prediction of IHI scores. A: Values of the adjusted coefficient of determination using from 1 to 40 eigenvectors resulting from the PCA. B: the coefficient correlation corresponding to the coefficient of determination of A. C: The *p*-values in −*log*_10_ of the corresponding coefficient of determination. D: Cross validation of the models using 20 to 40 dimensions by 100-fold. The red cross indicates the *MSE* of the model predicted using random values, and the errorbar corresponds to the standard deviation of *MSE* computed from 10,000 cross validations for each model, the triangle corresponds to the average *MSE*.

## 6 Discussion and Conclusion

In this paper, we proposed a method for template-based shape analysis using diffeomorphic centroids. This approach leads to a reasonable computation time making it applicable to large datasets. It was thoroughly evaluated on different datasets including a large population of 1000 subjects.

The results demonstrate that the method adequately captures the variability of the population of hippocampal shapes with a reasonable number of dimensions. In particular, Kernel PCA analysis showed that the large population of left hippocampi of young healthy subjects can be explained, for the metric we used, by a relatively small number of variables (around 50). Moreover, when a large enough number of subjects was considered, the number of dimensions was independent of the number of subjects.

The comparisons performed on the two small datasets show that the different centroids or variational templates lead to very similar results. This can be explained by the fact that in all cases the analysis is performed on the tangent space to the template, which correctly approximates the population in the shape space. Moreover, we showed that the different estimated centres are all close to the Fréchet mean of the population.

While all centres (centroids or variational templates) yield comparable results, they have different computation times. IC1 and PW centroids are the fastest approaches and can save between 70 and 90% of computation time over the variational template. Thus, for the study of hippocampal shape, IC1 or PW algorithms seem to be more adapted than IC2 or the variational template estimation. However, it is not clear whether the same conclusion would hold for more complex sets of anatomical structures, such as an exhaustive description of cortical sulci (40). Besides, one should note that, unlike with the variational template estimation, centroid computations do not directly provide transformations between the centroid and the population which must be computed afterwards to obtain momentum vectors. This requires *N* more matchings, which doubles the computation time. Even with this additional step, centroid-based shape analysis stills leads to a competitive computation time (about 26 hours for the complete procedure on the large dataset of 1000 subjects).

The iterative centroid estimation method can be used directly to estimate a template of a large population of shapes.

In future work, this approach could be improved by using a discrete parametrisation of the LDDMM framework (41), based on a finite set of control points. The control points number and position are independent from the shapes being deformed as they do not require to be aligned with the shapes’ vertices. Even if the method accepts any kind of topology, for more complexe and heavy meshes like the cortical surface (which can have more than 20000 vertices per subjects), we could also improve the method presented here by using a multiresolution approach (42). An other interesting point would be to study the impact of the choice of parameters on the number of dimensions needed to describe the variability population (in this study the parameters were selected to optimize the matchings). Finally we can note that this template-based shape analysis can be extended to data types such as images or curves.

## Acknowledgments

The research leading to these results has received funding from ANR (project HM-TC, grant number ANR-09-EMER-006, and project KaraMetria, grant number ANR-09-BLAN-0332), from the CATI Project (Fondation Plan Alzheimer) and from the program “ Investissements ďavenir” ANR-10-IAIHU-06.

IMAGEN was supported by the European Union-funded FP6 (LSHM-CT-2007-037286), the FP7 projects IMAGEMEND (602450) and MATRICS (603016), and the Innovative Medicine Initiative Project EU-AIMS (115300-2), Medical Research Council Programme Grant “Developmental pathways into adolescent substance abuse” (93558), the NIHR Biomedical Research Centre “Mental Health” as well as the Swedish funding agency FORMAS. Further support was provided by the Bundesministerium für Bildung und Forschung (eMED SysAlc; AERIAL; 1EV0711).

It should be noted that a part of this work has been presented for the first time in Claire Cury´s PhD thesis (43).

## Footnotes

1 http://www.imagen-europe.com/

1 http://fsl.fmrib.ox.ac.uk/fsl/fslwiki/FslOverview

1 http://www.brainvisa.info

